# *Galleria mellonella* model for studying Gram-positive bacterial implant biofilms

**DOI:** 10.1101/2025.02.18.638826

**Authors:** Kamran A. Mirza, Sandor Nietzsche, Tinatini Tchatchiashvili, Oliwia Makarewicz, Mathias W. Pletz, Lara Thieme

## Abstract

Implant-associated biofilm infections, particularly those caused by *Staphylococcus aureus* and *Enterococcus faecalis*, present significant challenges in clinical settings, often necessitating surgical removal. This study investigates the potential of the invertebrate model *Galleria mellonella* for evaluating biofilm formation of *S. aureus* and *E. faecalis* clinical isolates, the leading causes of implant-associated infections. Utilizing expanded polytetrafluoroethylene (ePTFE) sutures as a surrogate for cardiac implants, we employed two biofilm formation methodologies reflecting the two main routes of implant infections: *in vivo* biofilm development within larvae mimicking hematogenous spread and pre-formed biofilm transplantation mimicking contamination during surgery. Scanning electron microscopy revealed complex biofilm structures for the biofilm formed inside of the larvae on implant, closely mimicking clinical conditions. Antibiotic treatments with vancomycin and rifampicin demonstrated significant reductions in bacterial biofilms, proving highly effective. The study highlights the *G. mellonella* model’s potential for preclinical biofilm research, offering a cost-effective and ethical alternative to vertebrate models while providing valuable insights into biofilm-related infections and their treatment.

## 1 Introduction

Implant-associated biofilm infections present a significant challenge in medicine. Despite the combination of surgical debridement and antibiotic therapy, 30–40% of cases result in treatment failure, and necessitating invasive procedures like implant removal^1^. *Staphylococcus aureus* and *Enterococcus faecalis* are the primary pathogens implicated in these infections, with *S. aureus* being the leading pathogen in prosthetic valve endocarditis^2,3^. Both pathogens are well-known biofilm formers, contributing to the persistence of infective endocarditis on native and prosthetic valves and resistance to treatment^4,5^.As these infections persist, a more comprehensive understanding of biofilm formation on implants is crucial for developing effective strategies to overcome this enduring challenge.

The use of vertebrate models in studying implant-associated infections is limited by ethical concerns, particularly in experiments involving biofilm formation and treatment^6^. This is due to the prolonged observation and invasive procedures, significant stress, pain, and potential suffering for the animals involved. In response to these concerns, the 3R principles (replacement, reduction, refinement) have driven the search for alternative models^7^. The larvae of the greater wax moth aka *Galleria mellonella* have emerged as a promising, cost-effective, and ethically favourable model for studying human bacterial pathogens^8,9^. With an innate immune system similar to mammals and a proven susceptibility to diverse human pathogens, *G. mellonella* provides clinical relevant insights into implant-associated biofilm formation and treatment strategies.

In a previous study, implant biofilms were studied using a single bacterial strain of *S. aureus*^10^ and a nylon bristle to mimic an implant^11^. This study investigates implant-associated biofilm formation of three distinct clinical isolates of *S. aureus* and *E. faecalis* in the *G. mellonella* model. We introduced a expanded polytetrafluoroethylene (ePTFE) suture as a surrogate implant, mimicking prosthetic valve endocarditis pathogenesis, where biofilm formation starts on the suture material at the tissue-implant interface^12^. This research presents a comparative analysis of two biofilm formation methodologies – reflecting the two main routes of implant associated infections, i.e. *in vitro-*biofilm formation reflecting pathogen contamination during surgery (PF) and *in vivo* biofilm formation reflecting infection by hematogenous spreading pathogens (IL) within the larvae, using scanning electron microscopy to gain an in-depth understanding of biofilm structures. Finally, we assess the model’s efficacy in evaluating antimicrobial activity against bacterial biofilm by applying the recommended antibiotic regimen of vancomycin, rifampicin and their combination^13^ for *S. aureus* prosthetic endocarditis, examining its impact on biofilm formation and eradication.

## 2 Material and Methods

### 2.1 Bacterial isolates and implants

Clinical isolates of *S. aureus* (SA14073, SA29552, and SA31685) (Supplementary data Table S1) and *E. faecalis* (EF67230, EF1653, and EF9367) (Supplementary data Table S2) were previously collected in studies approved by the ethical committee of Jena University Hospital (3852/07-13 and 3694-02/13). The minimum inhibitory concentrations (MICs) of the antibiotics were assessed *via* VITEK2 (bioMérieux, Marcy-l’Étoile, France) and interpreted according to the clinical European Committee of Antimicrobial Susceptibility Testing (EUCAST) breakpoints in 2022. *S. aureus* isolates were cultured in Mueller Hinton (MH) broth (Oxoid Deutschland GmbH, Wiesel, Germany) and *E. faecalis* isolates were cultured in Todd-Hewitt (TH) broth (Oxoid Deutschland GmbH, Wiesel, Germany) at 37°C in a shaking incubator for 2 hours to reach the early exponential phase. The bacterial cultures were adjusted to an optical density at 600 nm (OD600) of 0.08 using a Multiscan GO spectrophotometer (Thermo Fisher Scientific, Waltham, USA), which corresponds to a bacterial count of approximately 10^8^ colony forming units (CFU) per mL and represented the working suspension for most experiments.

Two clinical implant materials were tested: the polyurethane (PU) intravenous catheter (Braun SE, Melsungen, Germany) and the polytetrafluoroethylene (ePTFE) cardiovascular suture (W. L. Gore & Associates GmbH, Putzbrunn, Germany). The PU catheter was sliced into fine bristles of approximately 0.2 mm (± 0.05 mm) in diameter and 0.5 cm in length, and ePTFE suture was cut in 0.5 cm in length (as it was already of 0.2 mm in diameter) using a sterile scalpel. The PU catheter was used only in the *in vitro* biofilm formation experiments to i) establish the experimental set up and to ii) compare the *in vitro* biofilm formation on both materials. The ePTFE sutures were than used to compare different methodologies of biofilm formation in the larvae.

### 2.2 *In vitro* biofilm formation on implants

Bacterial cultures at early logarithmic phase were adjusted to an optical density at 600 nm (OD600) of 0.08 using Multiscan GO spectrophotometer (Thermo Fisher Scientific, Waltham, USA) in a 1-cm cuvette and 200 µL/well was inoculated in microtiter plate in triplicates. The PU catheter was sliced into fine bristles of approximately 0.2 mm in diameter and 0.5 cm in length using a sterile scalpel and each slice was incubated at 37°C in a shaking incubator at 160 RPM in the bacterial suspension in the microtiter plate. The medium was refreshed after 24 hours of incubation. After 48 hours, bacterial suspension was removed, and the bristles were washed twice with 200 µL of 1 x PBS, sonicated for 15 minutes, vortexed for 30 seconds, serially diluted, plated on MH agar plates, and CFU was counted after 24 h of incubation at 37°C. The same setup was repeated with ePTFE suture.

### 2.3 Systemic infection in *G. mellonella* larvae

*G. mellonella* wax moth larvae were obtained from Bruno Mariani-FLOTEX (Augsburg, Germany). Larvae were stored in a refrigerator at 15 °C for a maximum of 2 weeks. In the experiments, larvae of an average size of 3.3 cm (with a standard deviation (SD) of ±0.12 cm) were used.

For induction of systemic infections in the larvae, early log phase bacterial cultures were centrifuged at 3000 g for 10 min, supernatants were removed, and the bacteria were resuspended in 1 x phosphate buffered saline (PBS, Carl Roth GmbH, Karlsruhe, Germany) to obtain 10^9^ CFU/mL. Bacterial suspensions of 10^9^ CFU/mL were serially 1:10 diluted and 10 µL of 10^9^, 10^8^ and 10^7^ CFU/mL were injected into the proleg of the larvae using a Hamilton syringe (Hamilton Bonaduz AG, Bonaduz, Switzerland) obtaining 10^7^, 10^6^ and 10^5^ CFU/larvae, respectively. Larvae survival was monitored every 24 h until 72 h and Kaplan-Meier curves were plotted. Each experiment contained five larvae per group and was repeated twice (n= 2×5= 10), including mock treatment (1 × PBS injection).

### 2.4 *In vivo* biofilm formation - establishing implant-associated infections within the larvae

Two methodologies were applied to compare the biofilm formation on both materials under different conditions: i) biofilm formation on the implant inside the larvae reflecting infection by hematogenous spreading pathogens (IL) and ii) pre-formed biofilm on the implant (PF) transplanted in the larvae reflecting pathogen contamination during surgery (Figure 1). For the larvae experiments, the PU catheter was ruled out because of the uneven surfaces due to manual cutting with a scalpel. The larvae experiments proceeded with ePTFE material with three clinical isolates of each bacterial species.

**Figure 1.**
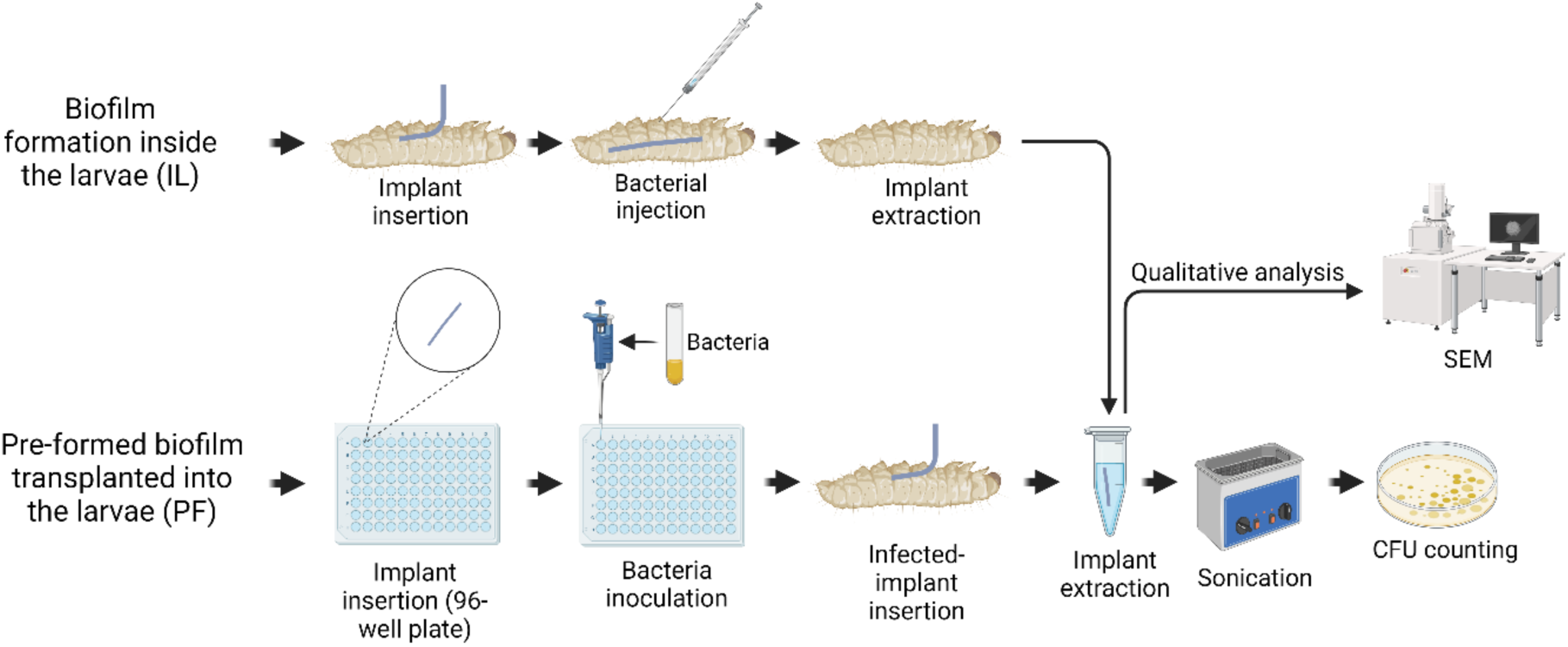
Schematic of biofilm generation methods and implementation of the material. IL = Biofilm formation on implants within larvae and PF = Transplantation of pre-formed biofilms on implants into larvae. Post-implantation survival was observed for 48 hours, after which implants were removed for CFU quantification and subjected to microscopic and microbial analyses.

**Table 1.**
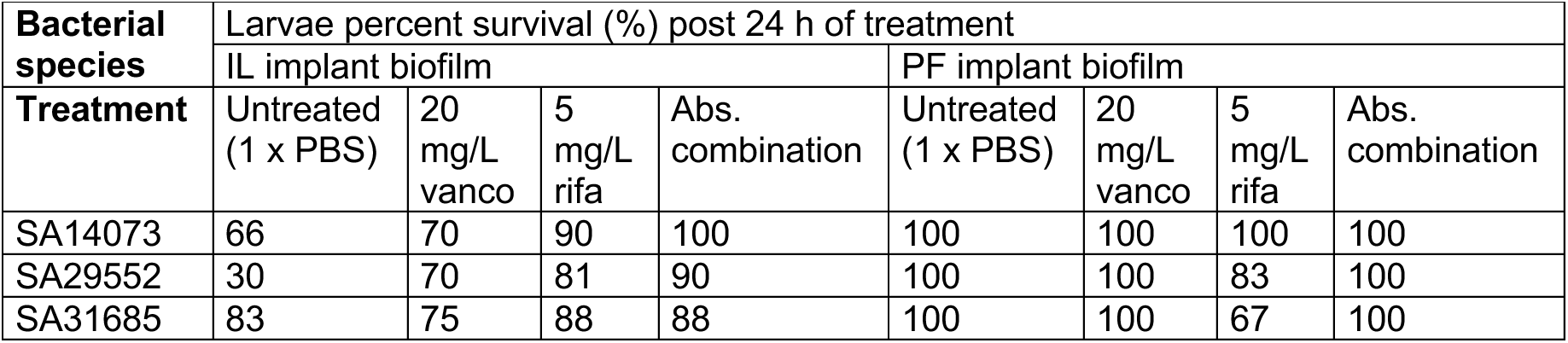
Percent survival (%) of the larvae after 24 h of treatment against biofilms formed by *S. aureus* isolates applying the IL or PF method.

In the first group (n=6 x 2=12), mimicking the hematogenous route of implant infection (IL), the implant was implanted 1 h before the larvae were infected with 10^5^ CFU/larvae through the proleg. In the second group (PF) (n=6×2=12), the implant was pre-incubated with a bacterial suspension of OD600 of 0.08 in a 96-well microtiter plate for 24 h at 37 °C in a shaking incubator with 160 rpm, followed by implantation into the larvae through the proleg. A third group (n=6×2=12) of larvae was inserted with sterile implants and 10 µl of 1 x PBS as mock bacterial treatment. All larvae were incubated with implants for 48 h. Survival of the larvae was monitored as well as bacterial attachment on the implants by re-implanting the biofilm implants followed by CFU determination as described above for *in vitro* biofilm formation. As selective solid media, mannitol salt agar (MSA) (Roth, Karlsruhe, Germany) was used for *S. aureus* and TH agar supplemented with 8 mg/L tetracycline (AppliChem GmbH, Darmstadt, Germany) for *E. faecalis* strains to exclude the larvae’s native *Enterococcus* microbiome. Freshly prepared plates and antibiotics were used for this purpose.

### 2.5 Scanning electron microscopy (SEM)

Biofilm-coated implants either created *in vitro* or in the larvae were fixed with freshly prepared modified Karnovsky fixative (4 % w/v paraformaldehyde, 2.5 % v/v glutaraldehyde in 0.1 M sodium cacodylate buffer, pH 7.4) for 1 h at room temperature. After washing 3 times for 15 min each with 0.1 M sodium cacodylate buffer (pH 7.4), the fibers were dehydrated in ascending ethanol concentrations (30, 50, 70, 90 and 100 %) for 15 min each. Next, the samples were critical-point dried using liquid CO2 and sputter coated with gold (thickness approx. 2 nm) using a CCU-010 sputter coater (Safematic GmbH, Zizers, Switzerland). Finally, the specimens were investigated with a field emission SEM LEO-1530 Gemini (Carl Zeiss NTS GmbH, Oberkochen, Germany). Triplicates of each bacterial isolate per implant were subjected to scanning electron microscopy (SEM).

### 2.6 Antimicrobial treatments in the larvae

To compare the antibiotic biofilm eradicating efficacy between the IL and PF routes of larval implant infection, larvae were treated with single and combined doses of vancomycin and rifampicin. For the PF experiment, the implants were immersed in a bacterial suspension adjusted to OD600 of 0.08 for 24 h and then transplanted into larvae. One hour after transplantation, larvae were treated with 10 µL of either 20 mg/L of vancomycin, 5 mg/L of rifampicin, or a combination of 20 mg/L vancomycin and 5 mg/L rifampicin (both Sigma-Aldrich Chemie GmbH, Taufkirchen), administered via a Hamilton syringe through a different proleg to prevent hemolymph loss. For the IL experiment, larvae were transplanted with sterile implants and post 1 h transplantation infected with 10^5^ CFU/larvae bacterial dose and incubated for 24 h at 37°C. Post 24 h incubation, alive larvae were treated with the same antibiotic dosages as described above. The concentrations were selected based on human blood peak plasma concentration of vancomycin^14^ and rifampicin^15^. The dosage was adjusted based on larvae hemolymph average volume of 50 µL. After 24 h of treatment, larvae were surgically opened, and the implants were carefully removed. The implants were washed twice with 1x PBS, sonicated for 15 minutes, and plated for CFU counts as described above.

## 3 Statistical analysis

All data were analyzed using GraphPad Prism 9 (GraphPad Software Inc., San Diego, USA). The Kaplan-Meier curves were compared using Log-rank (Mantel-Cox) test, CFU count of the bacterial attachment was compared using Two-Way ANOVA, and CFU reduction was compared using Kruskal-Wallis test. *P*-values of <0.05 were considered significant.

## 4 Results and Discussion

### 4.1 Bacterial biofilm *in vitro*

The PU and ePTFE materials were used as the artificial surfaces to quantify *in vitro* bacterial biofilm formation. Quantification of bacteria revealed an average of 10^6^ to 10^7^ CFU per implant on both ePTFE (Figure 2) and PU (Supplementary data Figure S1) across all bacterial isolates. In general, the variability in CFU counts among bacterial strains within a species was not high: the CFU count of the SA31685 (methicillin sensitive *S. aureus*, MSSA) isolate was approximately half a log10 higher compared to the SA14073 (methicillin resistant *S. aureus*, MRSA) isolate on the ePTFE implant. Similarly, EF67230 isolate showed 1 log10 higher CFU count compared to EF9367 isolate on the PU implant (Supplementary data Figure S1). This increase might be due to differences in cell surface properties or variations in the genetic regulation of adhesion factors^16^, as the statistical analysis demonstrated a significant difference in CFU counts among the isolates (Supplementary data Table 3). In summary, both *S. aureus* and *E. faecalis* exhibited substantial bacterial attachment to the implants in the *in vitro* setting.

**Figure 2.**
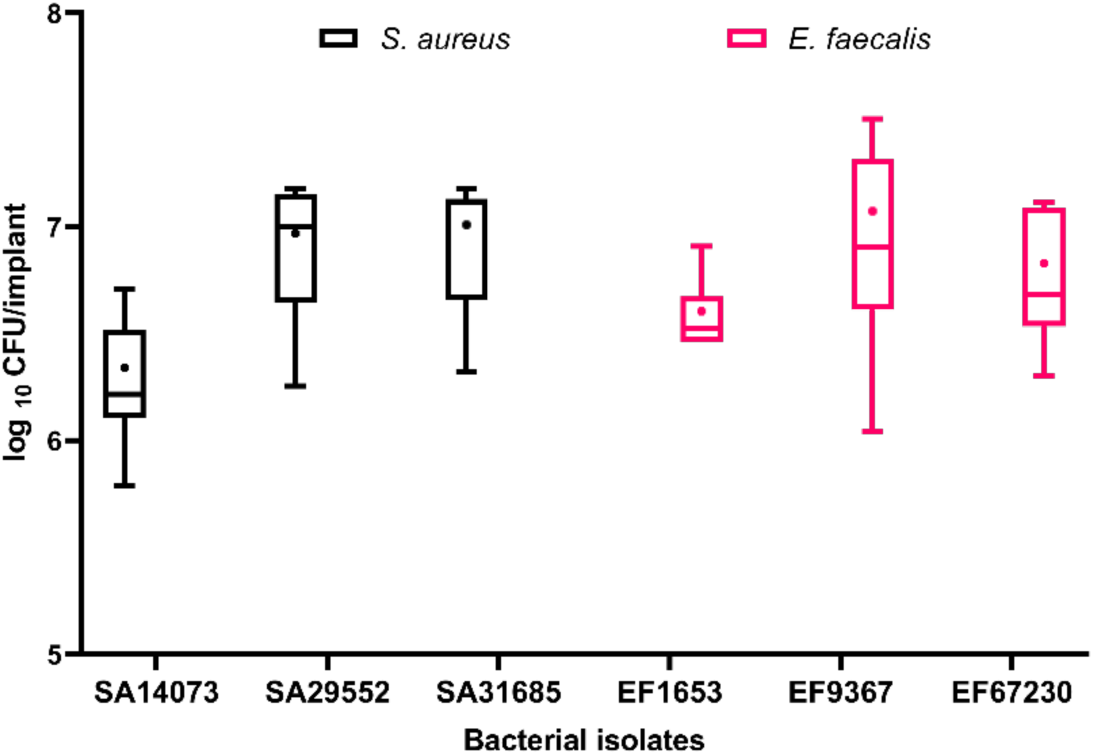
The quantification of *S. aureus* and *E. faecalis* isolates *in vitro* bacterial attachment on the ePTFE implant. The implants were incubated with bacterial suspension in 96 well plate for 48 h in a shaking incubator. After incubation, implants were washed twice with 1 x PBS and plated for CFU count. The experiments were conducted using triplicate implants per isolate and independently repeated twice (n= 3×2 = 6). The data was compared using One-Way ANOVA and P< 0.05 was considered significant (Supplementary data Table 3). The boxes and whiskers represent 95^th^-5^th^ percentiles, + represents the mean and the line represents the median.

In addition to bacterial quantification, the biofilms on the PU (Supplementary data Figure 2) and ePTFE (Figure 3) materials were visualized using SEM. The results depicted the ePTFE implants without any bacterial attachment that served as the control, whereas, the remaining SEM images displayed evident initial biofilm formation for both *S. aureus* and *E. faecalis* isolates (Figure 3). In the case of *S. aureus*, bacterial clusters with minimal extracellular matrix were observed, indicative of an early stage in biofilm formation (Figure 3 d-f). *E. faecalis* showed initial attachment and microcolony formation (Figure 3 g-i). Similarly, the *E. faecalis* isolate exhibited a sheet-like structure rather than clustered formations on the PU implant, consistent with the typical biofilm pattern exhibited by *E. faecalis* (Supplementary data Figure 2). In biofilm early stages, the first step is initial attachment (bacterial attachment to surface) followed by microcolony formation in which bacteria begin to multiply and form small aggregates with small production of biofilm matrix^17^.The absence of intricate or complex biofilm may be attributed to the absence of biological system, such as *in vivo* experimental conditions. The development of biofilm under *in vivo* conditions is facilitated by various protein and polysaccharides, ultimately contributing to the formation of sophisticated biofilm structures^18^. To summarize, SEM analysis revealed an early stage of biofilm development and indicated the use of *in vivo* system.

**Figure 3.**
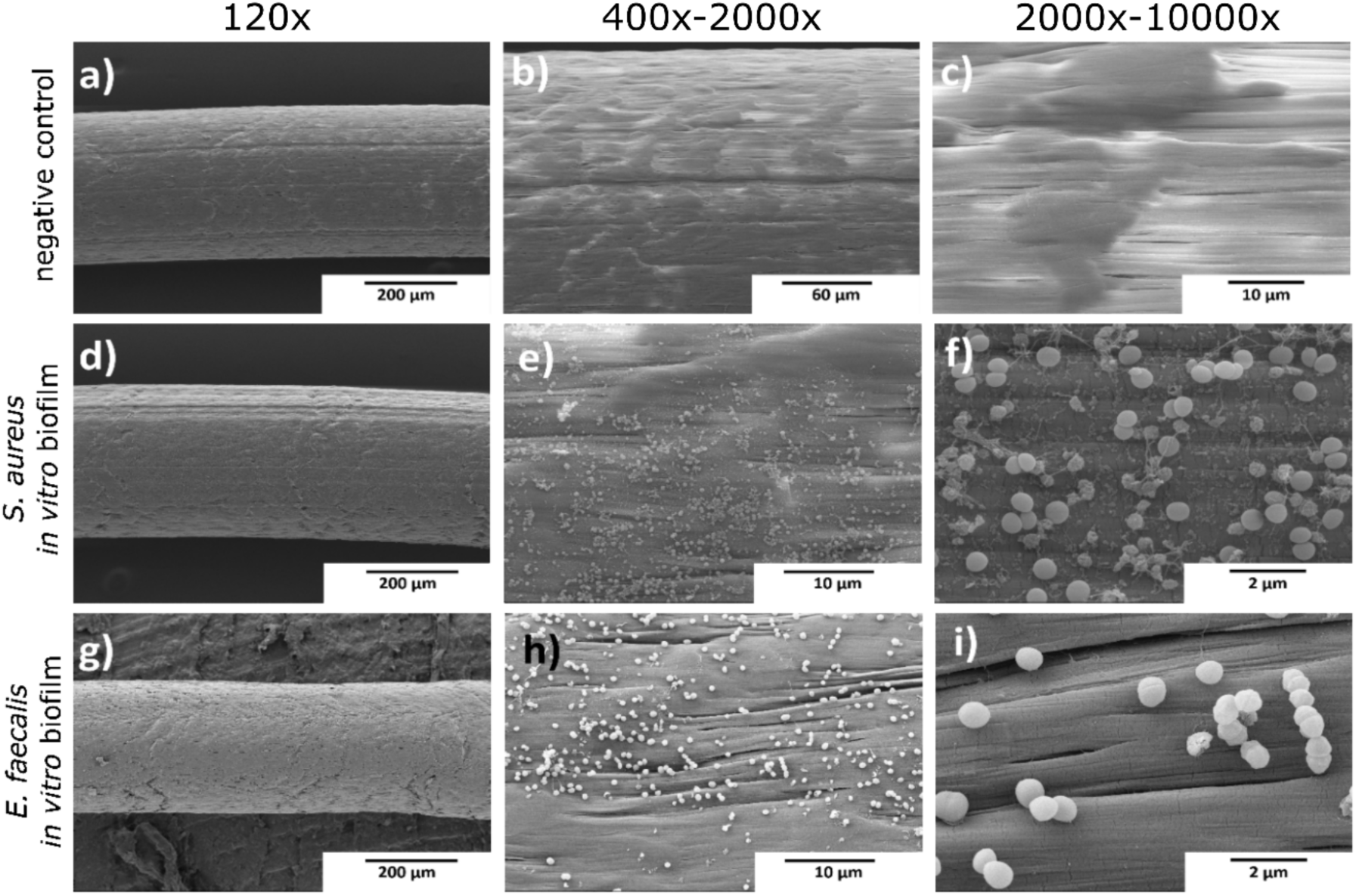
SEM images of ePTFE implant without bacterial attachment (a to c) and *in vitro* colonisation by *S. aureus* isolate SA14073 (d-f) and *E. faecalis* isolate EF1653 (g-i) for 48 h. The images showed different magnifications as indicated. The scale bar (black) represents 200 µm (a, d, g), 60 µm (b), 10 µm (c, e, h,), and 2 µm (f, i).

### 4.2 *G. mellonella* and systemic infection

To mimic implant-associated biofilm infections in *G. mellonella* larvae, determining the optimal bacterial infection dosage was essential. Upon injection, the larvae’s immune response is characterized by pathogen encapsulation and melanin production, which manifests as visible darkening of the larvae^19^. This pigmentation change served as a practical indicator to assess the impact of varying bacterial dosages on larval health. It was imperative to balance promoting biofilm formation while preventing systemic infection that could result in larval mortality. Therefore, larvae were administered a series of 10-fold increasing bacterial dosages, followed by a 72 h incubation period, with survival monitored at 24 h intervals. The results revealed a significant correlation between the inoculation dosage of bacterial isolates and larval mortality rates (Supplementary data Table 3). Specifically, an inoculum of 10^5^ CFU demonstrated a larval survival rate exceeding 80% after 72 h of infection (Figure a-c). Conversely, higher bacterial dosages, such as 10^7^ CFU, resulted in 100% larval mortality (Figure 4a). Additionally, with an inoculum of 10^6^ CFU, the larval survival rate remained below 80% (Figure 4 a, b). A similar pattern was observed in larvae infected with *E. faecalis* isolates, where increasing bacterial dosages corresponded with higher larval mortality rates (Figure 4 d, e). Notably, an exception was observed with *E. faecalis* isolate 67230, where even a lower dosage of 10^5^ CFU led to a significant reduction in survival, with less than 50% of the larvae surviving after 72 h of infection, indicating the high virulence of this isolate (Figure 4 f). The observed variations in pathogenicity among bacterial strains can be attributed to several factors, including differences in the host’s immune response and the genetic makeup of the bacteria. The innate immune variability among larvae contributes to differing levels of susceptibility to specific pathogens. Moreover, certain bacterial strains have evolved mechanisms that enhance their pathogenicity towards specific hosts^20^. For example, a study utilizing the *G. mellonella* infection model with *E. faecalis* reported rapid larval death within 30 minutes post-infection with 10^5^ CFU, attributed to the activity of gelatinase, an enzyme that degrades inducible antimicrobial peptides in the larvae^21^. Consequently, based on these findings, a bacterial dosage of 10^5^ CFU per larva was selected for subsequent methodologies concerning bacterial biofilm formation on implants within the larvae.

**Figure 4.**
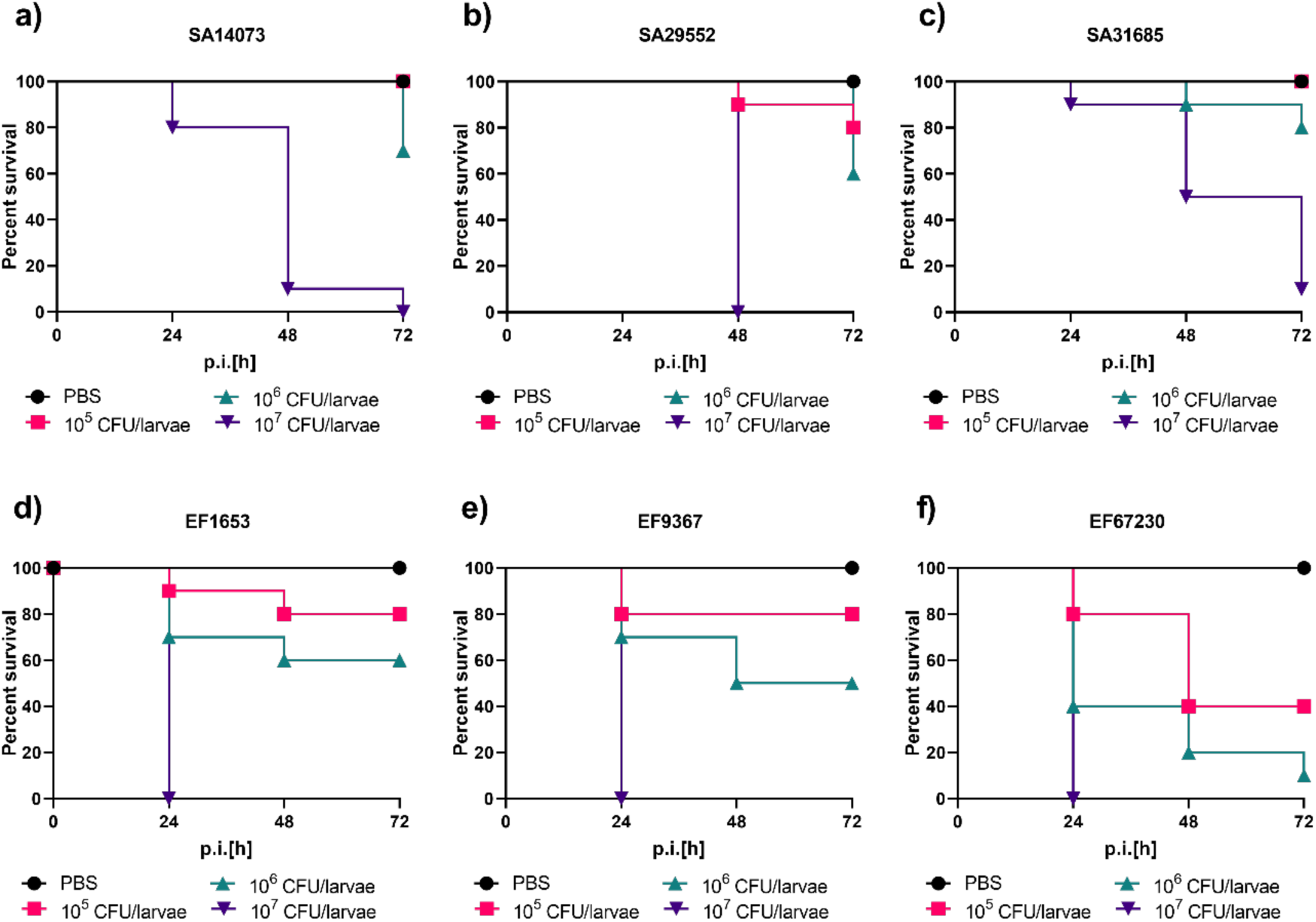
Survival curves of the larvae infected with different dosage of *S. aureus* isolates SA14073 (a), SA29552 (b) and SA31685 (c) and *E. faecalis* isolates EF1653 (d), EF9367 (e) and EF67230 (f). The larvae were infected with different bacterial dosages via proleg, followed by a 72 h incubation period, with survival assessments conducted every 24 h. Five larvae per bacterial dose were tested and the experiment was repeated twice (n= 5×2= 10). The statistical significances were analysed using Log-Rank test and mentioned in Supplementary data Table 3.

### 4.3 Comparison of different implant biofilm methodologies in *G. mellonella*

To investigate the differences in biofilm formation and their interactions with *G. mellonella* larvae, we utilized two distinct methods: IL and PF. Larval survival was monitored every 24 hours. The results indicated that IL biofilm formation led to higher mortality across all bacterial isolates compared to PF biofilms (Fig. 5 a-b, Supplementary data Table 3). For example, larvae infected with *S. aureus* isolate 29552 showed a 33% survival rate with the IL method, while the PF method increased survival to 67% (Figure 5 a). Similarly, *E. faecalis* isolate 9367 resulted in an 8% survival rate with IL, compared to 92% with PF (Figure 5 b). The IL method, involving direct bacterial injection into the larval hemolymph, caused systemic infection and higher mortality. Additionally, larvae with pre-infected implants exhibited lower survival compared to those with mock treatment or planktonic infections, likely due to intensified immune responses and melanization triggered by the foreign implant^22^. In the PF biofilm, bacteria evade the larval immune system by forming a biofilm that is quickly encapsulated by host material, focusing on survival through evasion rather than direct confrontation^23^. In contrast, the IL biofilm requires bacteria to first defend against and attack the larval immune system before forming a biofilm on the implant, necessitating an aggressive “attack” phenotype^24^. In summary, PF biofilms on implants, transplanted into larvae, result in higher survival rates compared to biofilms formed directly within the larvae.

**Figure 5.**
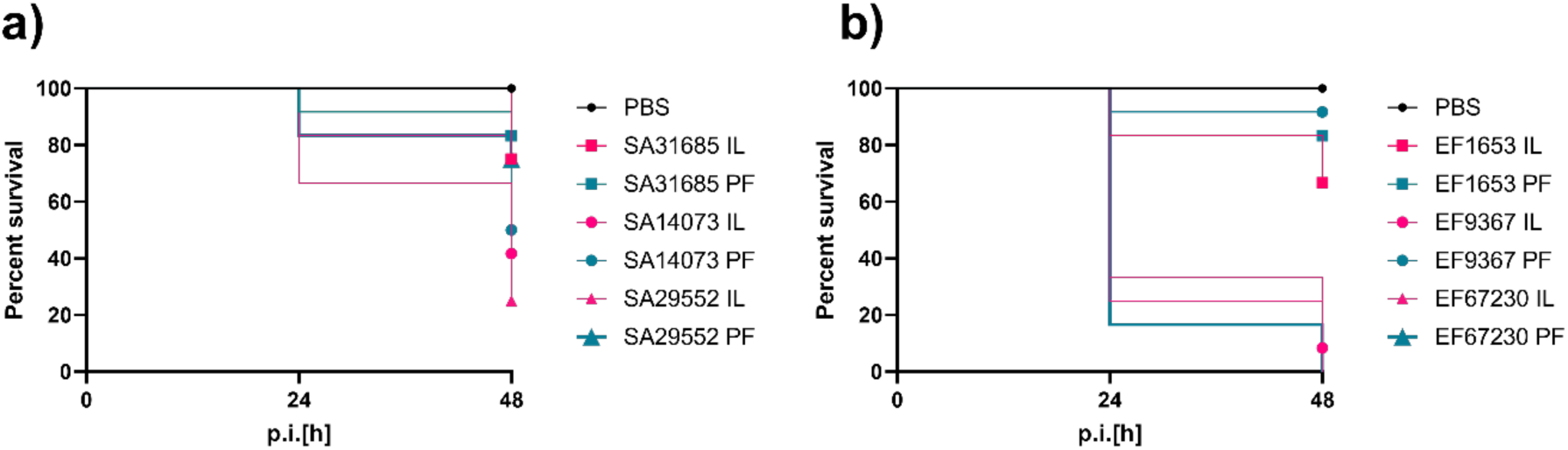
Survival of the larvae infected with implant biofilm of different methodologies IL and PF of *S. aureus* isolates (a) and *E. faecalis* isolates (b). Two distinct approaches were employed: IL biofilm, where the biofilm developed on the implant inside the larvae; and PF biofilm, involving biofilm formation on the implant *in vitro* prior to transplantation into the larvae. The larvae were incubated for 48 h and every 24 h survival was observed. The experiment utilized six larvae per bacterial isolate and was replicated twice (n= 6×2 = 12). The statistical differences were analyzed using Log-rank test and were mentioned in Supplementary data Table 3.

A quantitative analysis of biofilm formation on the implants was performed using CFU counts. The results indicated an average CFU range of 10^6^ to 10^7^ per implant for both the IL and PF methodologies (Figure 6 a-b), which closely matches the outcomes observed in the *in vitro* setting (Figure 2). This consistency in CFU counts between *in vivo* and *in vitro* conditions underscores the reproducibility of the biofilm quantification methods employed. A significant difference was observed with the *S. aureus* isolate 31685, a methicillin-sensitive strain (MSSA) (Figure 6 a). The IL biofilm exhibited a mean CFU count of 10^7^ per implant, compared to 10^6^ for the PF biofilms. MSSA strains often demonstrate robust biofilm formation capabilities, particularly in the presence of host factors such as fibrinogen and fibronectin, which are abundant in the host environment^25^. The IL method, which involves direct interaction with the host’s immune system and biological milieu, might provide these factors, thereby enhancing biofilm formation. The enriched nutrient availability and the supportive environment within the host may further contribute to the observed increase in biofilm biomass^11^.

**Figure 6.**
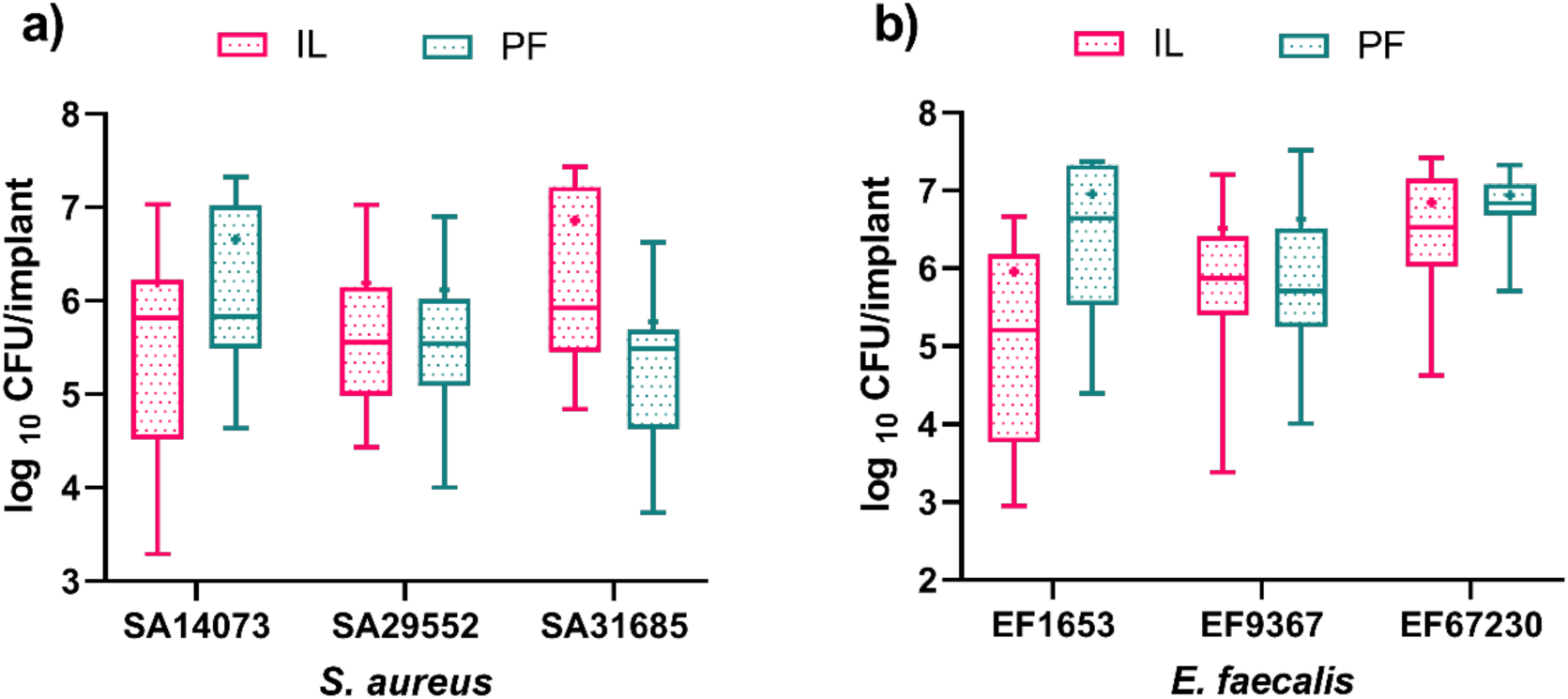
The CFU quantification of *S. aureus* isolates (a) and *E. faecalis* isolate (b) on the ePTFE implant. Two distinct methodologies were compared such as IL biofilms and PF biofilms as described above. Following a 48 h incubation period, the implants were removed and plated for CFU count. Six implants per bacterial isolate were included, and the experiment was conducted twice (n= 6×2 = 12). The data was represented as boxes and whiskers with 95^th^-5^th^ percentiles, + represents mean and the line represents the median. The data was compared using the Kruskal-Wallis test and significance difference was mentioned in Supplementary data Table 3.

After CFU quantification, SEM was used to visually compare biofilm formation on ePTFE implants between the IL and PF methods for *S. aureus* and *E. faecalis* isolates. The IL biofilms exhibited a notably complex structure, with SEM revealing complex biofilm formations predominantly on larval-derived material coating the implants (Figure 7). In contrast, sterile control implants showed only an overlay of larval material (Figure 7 a-c). SEM images of SA14073 (Figure 7 d-f) and EF1653 (Figure 7 g-i) demonstrated mature biofilms, with bacterial clusters embedded in an extracellular polymeric substance (EPS) matrix, largely derived from the host’s bodily material. This indicates that in implant-associated infections, bacteria often rely on host proteins for attachment and biofilm development, reducing the need to produce their own extracellular matrix^18^.

**Figure 7.**
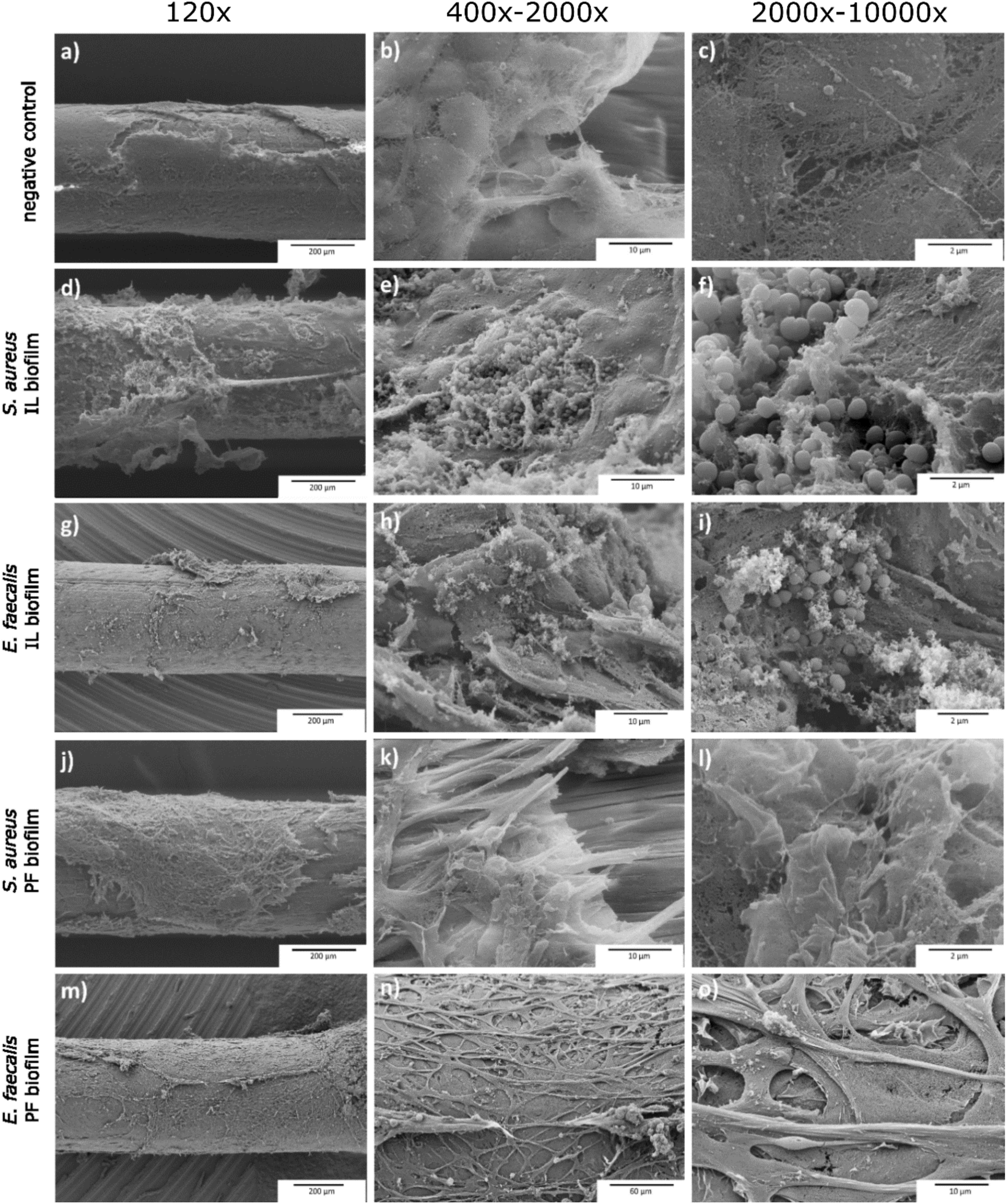
SEM comparison of IL and PF biofilm formation methods. Panels (a-c) display the SEM images of sterile ePTFE implants. Panels (d-f) and (g-i) illustrate biofilm formation by *S. aureus* isolate SA14073 and *E. faecalis* EF1653, respectively, using the IL method within *G. mellonella* larvae, where biofilm formation is facilitated by larval body materials. The sutures were inserted into the larvae through the proleg, and 1 h later, a bacterial dosage was injected. After incubation, the implants were removed and fixed for SEM analysis. Panels (j-l) and (m-o) show the SEM images of SA14073 and EF 1653, respectively using the PF method, indicating a noticeable absence of biofilm formation. The ePTFE suture was initially incubated with a bacterial suspension *in vitro* and subsequently transplanted into larvae *in vivo*. The IL method demonstrates complex biofilm development assisted by the larvae’s body material, while the PF method captures bacterial attachment encapsulated by larval body materials but lacks visualization of significant biofilm structure. This encapsulation suggests comparable CFU counts between both methods, yet highlights differences in biofilm development dynamics. The images showed different magnifications as indicated on the panels. The scale bar (black) represents 200 µm (a, d, g, j, m), 10 µm (b, e, h, k), 60 µm (n), and 2 µm (c, f, i, l) and 10 µm (o).

These findings underscore the key differences between *in vitro* (Figure 3) and IL (Figure 7 d-f) biofilm formation, with IL biofilms displaying a more complex and mature structure. This highlights the necessity of using *in vivo* models to study biofilms, particularly for determining Minimum Biofilm Eradication Concentrations (MBECs), as the complexities of *in vivo* biofilms significantly influence treatment efficacy.

In contrast to the complex IL biofilm formation, SEM analysis revealed no visible bacterial biofilm for both *S*A14073 (Figure 7 j-l) and EF1653 (Figure 7 m-o) isolates on the PF implants. Unlike the IL approach, where biofilm formation within the larvae utilized larval body materials (Figure 7 d-i), the PF implant showed bacterial attachment covered by larval body material. The CFU counts indicated a comparable level of viable bacterial attachment between both IL and PF biofilm formation methods (Figure 6). The observation at higher magnifications demonstrated that larval body material coated the implant (Figure 7 j-o), though no visible biofilm was observed. PF implants exhibited higher larval survival rates (Figure 5), likely due to the bacterial attachment being confined within the larval body material, which limited the bacterial spread. This pattern mirrors chronic infections, where bacteria persist in biofilms resulting in prolonged and recurrent infections. These findings highlight the differences between the PF and IL biofilm formation methodologies. The IL method exhibited complex biofilm development assisted by larval body materials, while the PF method showed encapsulated bacterial attachment. These distinctions suggest that the larval body material influences biofilm development dynamics and may influence the treatment outcomes.

### 4.4 Assessment of antimicrobial effects in *G. mellonella*

Effective evaluation of antimicrobial treatments against biofilm-associated implant infections necessitates a reliable *in vivo* model that accurately mimics clinical conditions. The *G. mellonella* larvae model offers a cost-effective and ethically favourable alternative for assessing antimicrobial effects. In this study, we evaluated the efficacy of commonly used antibiotics vancomycin and rifampicin, both alone and in combination, against *S. aureus* biofilm infections established in *G. mellonella* by evaluating survival rates, CFU reductions on the implants and visible changes in SEM images Both routes of implant-associated biofilm establishment (IL vs PF) were compared to analyse potential effects on the treatment outcome.

Our findings indicated that the combination of vancomycin and rifampicin combination or rifampicin alone significantly enhanced larval survival in biofilm-associated infections. Specifically, larvae infected with SA14073 IL (implant-in-larvae) biofilm exhibited a 100% survival rate following combination therapy, compared to 90% with rifampicin alone and 66% with the control treatment. Similarly, for the *S. aureus* 29552 IL biofilm, survival rates were 90% with combination therapy and 80% with rifampicin alone, compared to 30% for the control group. These results underscore the significant efficacy of rifampicin alone and in combination in biofilm eradication across multiple *S. aureus* isolates (Supplementary data Table 3).

However, the PF (pre-formed) biofilm model did not show a significant difference in larval survival with or without antibiotic treatment, suggesting its limited effectiveness in assessing survival outcomes post-treatment. The only exception was observed with rifampicin alone, which supported only 65% survival in larvae infected with SA31695, while the combination treatment led to 100% survival. This underscores a limitation of the PF model in evaluating antimicrobial efficacy *in vivo*, particularly when considering survival as an endpoint parameter.

### 4.5 CFU Quantification and SEM Analysis

CFU counts revealed significant reductions in bacterial attachment with both the combination treatment and rifampicin alone. For SA14073 IL biofilm, the combination treatment achieved a 2-log10 reduction, while rifampicin alone resulted in a 2-log10 reduction as well. Similar reductions were observed with isolates SA29552 and SA31685 (Figure 8 a). In the PF biofilms, the antibiotic combination led to an even greater 5-log10 reduction in the SA14073 and SA29552 isolates, with rifampicin alone yielding a 5-log10 reduction (Figure 8 a-b). Both the combination therapy and rifampicin alone achieved a 4-log10 reduction for isolate SA31685 in the PF biofilms (Figure 8 c). These results were statistically significant, as detailed in Supplementary Table 3.

**Figure. 8.**
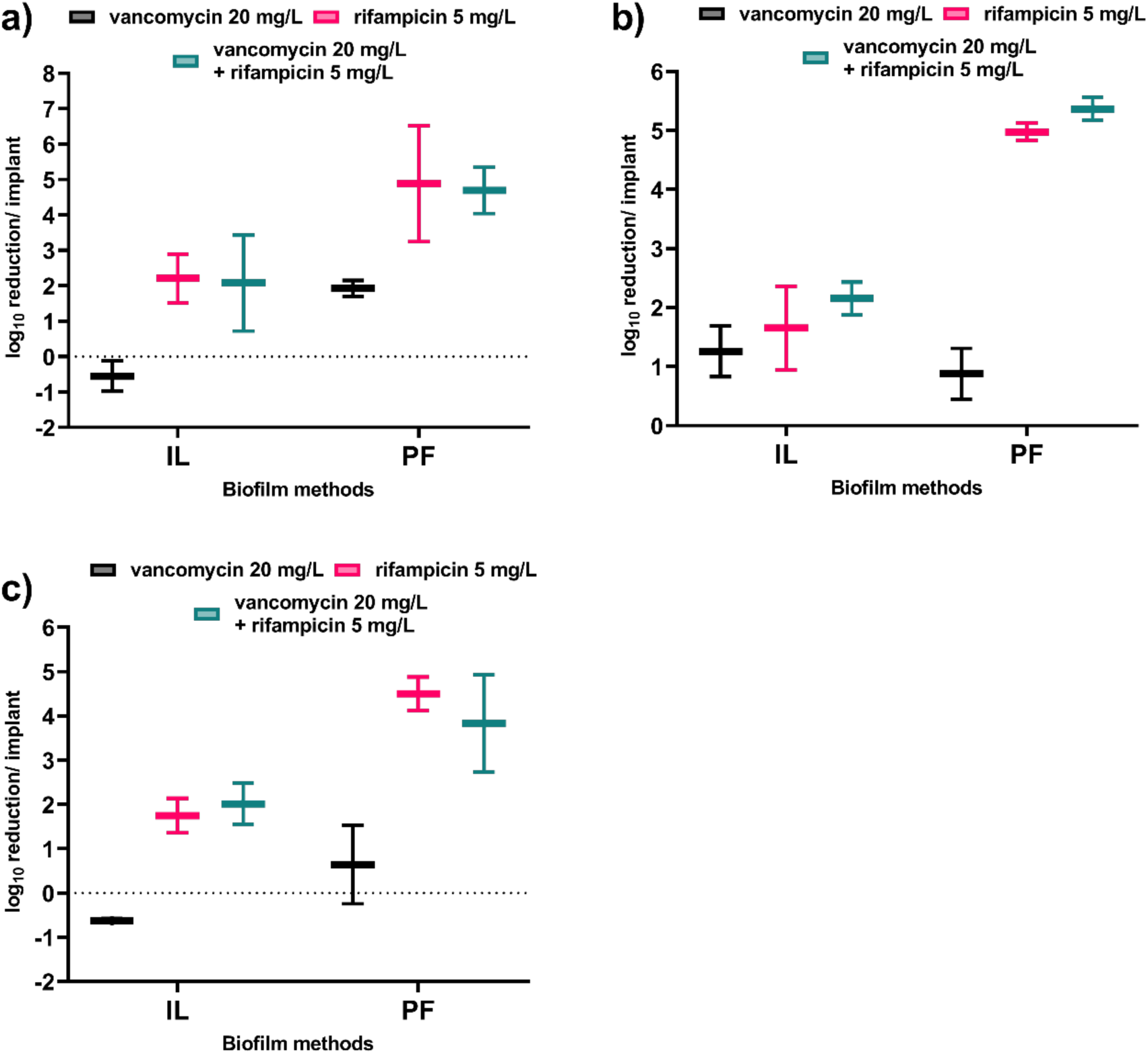
The log10 CFU reduction was calculated compared to untreated control to assess reduction of bacterial biofilm of *S. aureus* isolates SA14073 (a), SA29552 (b) and SA31685 on the implanted material by the commonly used antibiotics for prosthetic endocarditis. For the IL biofilm, larvae were infected post 1 h of implant insertion, incubated for 24 h, treated with antibiotics and incubated. Post 24 h of incubation, implant were quantified via CFU count. For the PF biofilm, the implants were pre-infected *in vitro* and subsequently transplanted into larvae. 1 h post-transplantation, larvae were subjected to injections of either vancomycin (20 mg/L), rifampicin (5 mg/L), or a combination of vancomycin (20 mg/L) and rifampicin (5 mg/L). Three larvae per bacterial isolate were included in the experiment, which was replicated twice independently (n= 3×2= 6). The data was compared using Kruskal-Wallis test and P<0.05 was considered significant. The data was represented as boxes and whiskers with min to max, the line represents the mean and significance difference was mentioned in Supplementary data Table 3.

The efficacy of the combination of vancomycin and rifampicin has been debated in previous studies, with some suggesting a synergistic effect, while others found rifampicin alone to be highly effective in models such as zebrafish^21^. In our study, no synergistic effect was observed; rifampicin alone was sufficient to significantly improve larval survival. Interestingly, the higher reductions in CFU counts observed in PF biofilms may be attributed to the encapsulation of the biofilm by larval body material. This encapsulation likely restricts bacterial dissemination, thereby enhancing the effectiveness of the rifampicin alone or in combination.

SEM imaging, whether from the IL method (Figure 9) or the PF method (Supplementary data Figure 3), showed a notable absence of bacteria after treatment with vancomycin, rifampicin, or their combination (untreated controls showed in Figure 7 d-i). At higher magnifications, damaged bacterial cells were evident across all treatment groups, reflecting the expected action of vancomycin and rifampicin. Vancomycin, which targets cell wall synthesis, effectively lysed bacterial cells on the surface. However, its larger molecular size likely limited its penetration into deeper tissue and biofilm layers^14^ within the larvae, reducing its overall effectiveness against bacteria embedded in the larval body material. Additionally, vancomycin is less effective against persister cells, as it primarily targets replicating bacteria through cell wall synthesis inhibition. Although these deeper biofilms were not detectable by SEM, their viability was confirmed through CFU counts.

**Figure 9.**
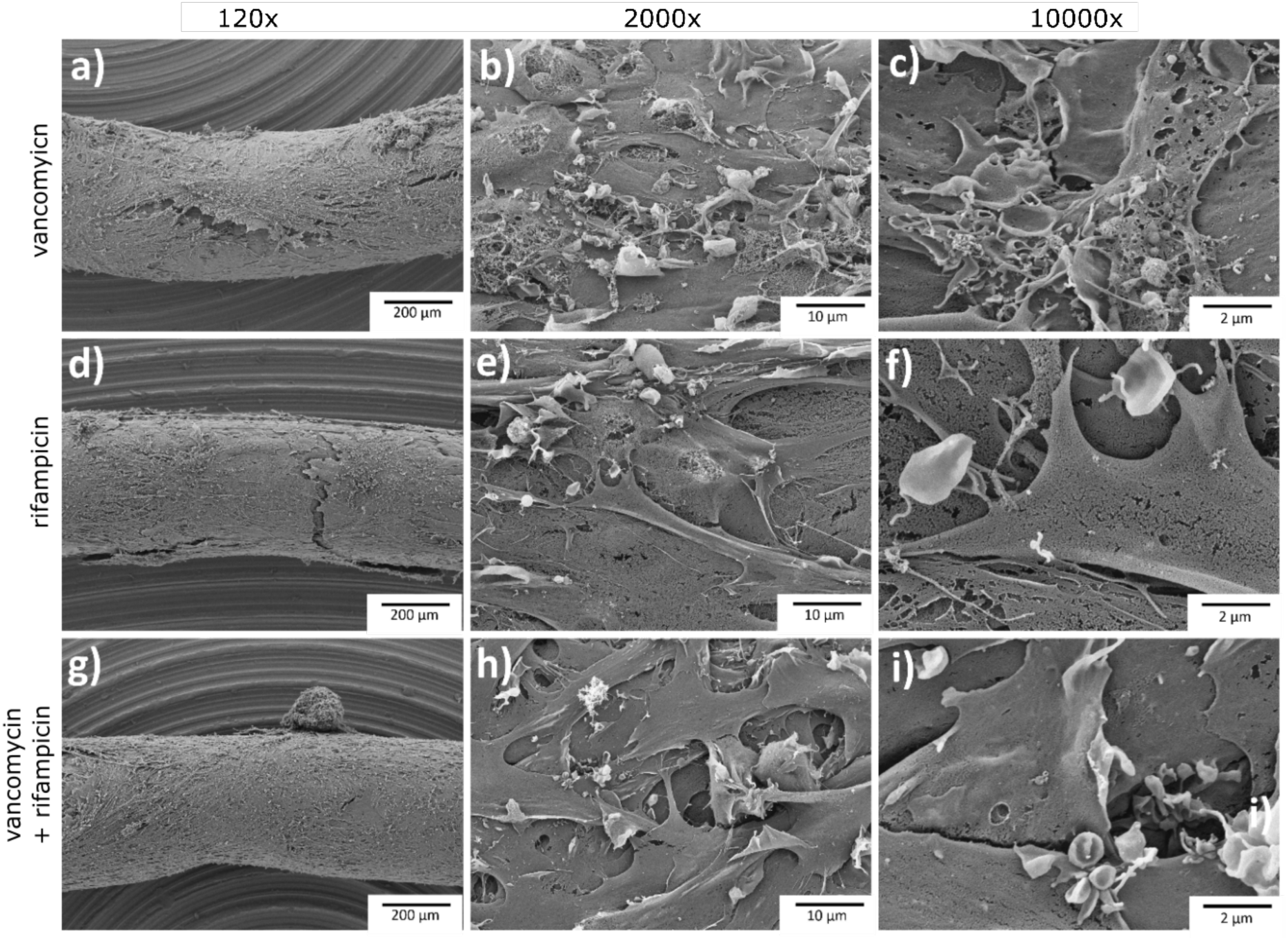
The post antibiotic treatment SEM analysis of *S. aureus* isolate SA14073 biofilms formed on implants with IL methodology. The biofilm treated with 20 mg/L vancomycin (a-c), 5 mg/L rifampicin (d-f) and combination of vancomycin and rifampicin (g-i). The larvae were injected with SA14073 post 1 h insertion of the implant and incubated, post 24 h incubation, larvae were treated with antibiotics. After 24 h of treatment, implants were taken out of the larvae, washed with 1 x PBS, fixed and SEM was performed. The images showed different magnifications as indicated on the panels. The scale bar (black) represents 200 µm (a, d, g), 10 µm (b, e, h) and 2 µm (c, f, i) magnification.

Conversely, rifampicin, known for its superior tissue and biofilm penetration and effective against persister cells^26^, resulted in significant bacterial reduction, as corroborated by CFU data. This underscores a key limitation of SEM: its inability to penetrate or visualize biofilms encased within host body materials. To gain a more comprehensive understanding of biofilm persistence and eradication following antibiotic treatment, future studies should consider employing complementary imaging techniques.

IL biofilms more closely replicate the complexities of clinical implant-associated biofilm infections and are superior for survival monitoring and SEM imaging, whereas PF biofilms are less effective in this context. In our study, vancomycin alone did not significantly reduce CFU counts, likely due to its limited tissue and biofilm penetration, as well as its slow bactericidal activity—factors that necessitate its combination with other antibiotics in clinical settings. Rifampicin, on the other hand, demonstrated high tissue and biofilm penetration but is also known to promote bacterial resistance. The combination of vancomycin with rifampicin can enhance treatment efficacy and mitigate resistance risks; however, a study has shown that it does not completely prevent resistance development^27^

It is important to note that this study did not assess post-treatment bacterial resistance, as the primary focus was on validating the model. Future research should explore optimal treatment strategies for *S. aureus* biofilm infections, including a deeper investigation into the dynamics of antibiotic resistance following treatment.

## 5 Conclusion

This study investigates various methodologies for studying implant-associated biofilms using the *G. mellonella* invertebrate model. Expanded PTFE suture, commonly used in cardiac surgeries, was selected as the implant material to induce biofilm formation, with a focus on evaluating treatments for prosthetic endocarditis. The study uniquely addresses three clinical strains each of *S. aureus* and *E. faecalis*, marking a novel approach within this model. Multiple methods for inducing biofilm formation were applied, and electron microscopy was used to visualize biofilms and identify critical differences in their formation.

*In vivo* models are essential for understanding biofilm formation as they replicate conditions similar to difficult-to-treat infections. The *G. mellonella* model shows significant potential for proof-of-concept studies on implant-associated biofilms, offering cost-effectiveness, efficiency, and extensive data generation compared to vertebrate models. However, limitations include the lack of an adaptive immune system, and the absence of an organ like a heart in direct contact with the implant, restricting some conclusions. Despite these limitations, the model is promising for reducing reliance on vertebrate models and enhancing our understanding of biofilm-related infections.

## 8 Declaration

The authors declare no conflicts of interest. All data generated in the study are included in the manuscript and can be passed on to interested parties upon request.

## 9 Author contributions

Conceptualization: K.M., O.M., L.T.; Experimentation: K.M.; Formal analysis and investigation: K.M., O.M., L.T., S.N.; Writing-original draft preparation: K.M.; Writing-review and editing: K.M., T.T., O.M., L.T., M.P.; Supervision: O.M., L.T., M.P.

## 10 Data availability

The data supporting the findings of this study are available upon request. For access, please contact the corresponding author, Kamran Mirza.

