## Supplementary data for "*Galleria mellonella* model for studying Gram-positive bacterial implant biofilms"

**Table S1** Antibiogram of *S. aureus* strains used in this study.

| | ***S. aureus*** | | |
|---|---|---|---|
| **Strain** | **SA14073** | **SA29552** | **SA31685** |
| Specimen | Blood culture | Catheter material | Wound swab |
| Clinical background | Endocarditis | Catheter-related blood-stream infection | Wound infection |
| Macrolide | 8 (R) | 8 (R) | 0.25 (S) |
| Clindamycin | 8 (R) | 8 (R) | 0.25 (S) |
| Vancomycin | 1 (S) | 2 (S) | 1 (S) |
| Penicillin | 0.5 (R) | 0.5 (R) | 0.5 (R) |
| Flucloxacillin | 4 (R) | 4 (R) | 0.25 (S) |
| Linezolid | 2 (S) | 2 (S) | 2 (S) |
| Rifampicin | 0.5 (S) | 0.5 (S) | 0.5 (S) |
| Daptomycin | 0.25 (S) | 2 (R) | 0.25 (S) |
| Fosfomycin | 8 (S) | 8 (S) | 8 (S) |
| Ciprofloxacin | 8 (R) | 8 (R) | 0.5 (S) |
| Moxifloxacin | 4 (R) | 8 (R) | 0.25 (S) |
| Gentamicin | 0.5 (S) | 0.5 (S) | 0.5 (S) |
| Tigecycline | 0.12 (S) | 0.12 (S) | 0.12 (S) |
| Profile | MRSA | MRSA | MSSA |

* MRSA (methicillin-resistant *Staphylococcus aureus*), MSSA (methicillin susceptible *Staphylococcus aureus*), S (sensitive), R (resistant)

**Table S2** Antibiogram of *E. faecalis* strains used in this study.

| | ***E. faecalis*** | | |
|---|---|---|---|
| **Strain** | **EF67230** | **EF1653** | **EF9367** |
| Specimen | Mitral valve swab | Blood culture | Blood culture |
| Clinical background | Endocarditis | Urosepsis | Recurrent bacteremia |
| Vancomycin | 2 (S) | 1 (S) | 1 (S) |
| Teicoplanin | <0.5 (S) | < 0.5 (S) | <0.5 (S) |
| Linezolid | 2 (S) | 2 (S) | 2 (S) |
| Amoxicillin | <2 (S) | <2 (S) | 2 (S) |
| Ampicillin | <2 (S) | <2 (S) | 2 (S) |
| Moxifloxacin | 0.5 (S) | 0.5 (S) | 1 (S) |
| Levofloxacin | 2 (I) | <10 (R) | 2 (I) |
| Tigecycline | <0.12 S | <0.12 (S) | <0.12 (S) |

S (sensitive), R (resistant)

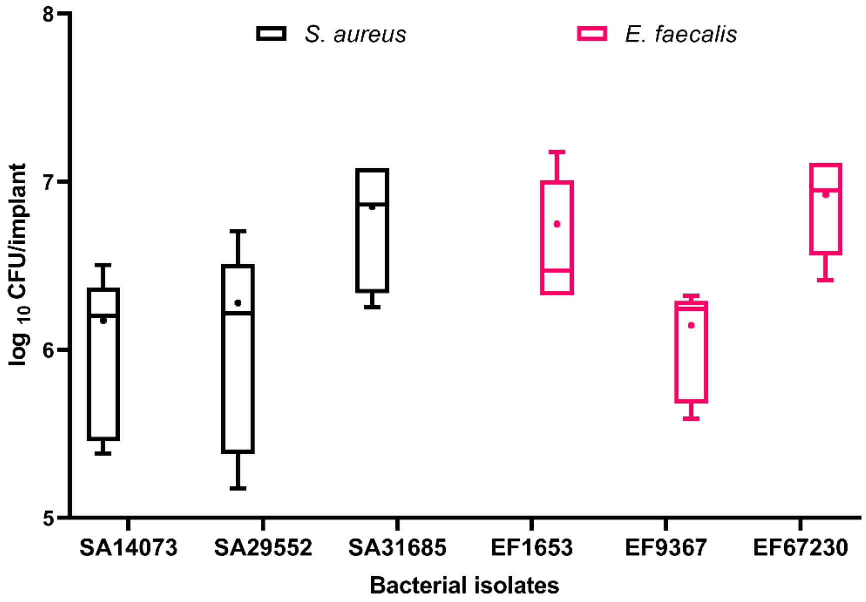


Figure S1. The quantification of *S. aureus* and *E. faecalis* isolates *in vitro* bacterial attachment on the PU implant. The implants were incubated with bacterial suspension in 96 well plate for 48 h in a shaking incubator. After incubation, implants were washed twice with 1 x PBS and plated for CFU count. The experiments were conducted using triplicate implants per isolate and independently repeated twice (n= 3x2 = 6). The data was compared using One-Way ANOVA and P< 0.05 was considered significant (Supplementary data Table 3). The boxes and whiskers represent 95th-5th percentiles, + represents the mean and the line represents the median

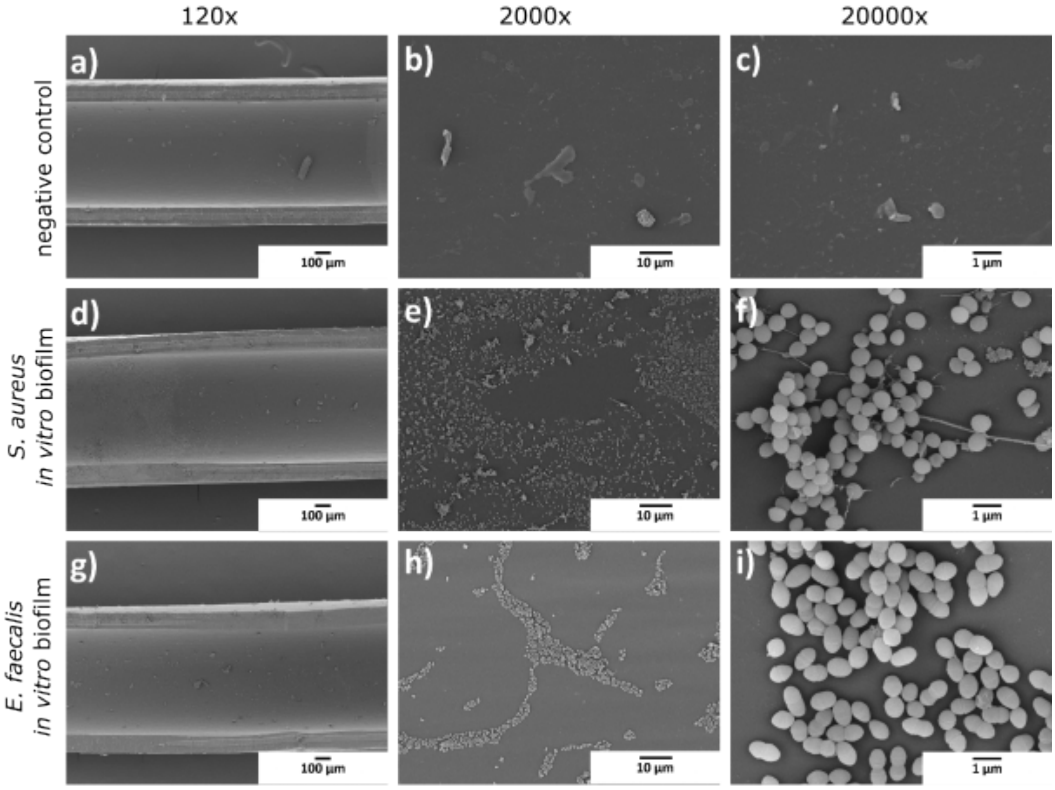


Figure S2. The SEM of PU implant without bacterial attachment (a to c) and *in vitro* colonisation by *S. aureus* isolate SA14073 (d-f) and *E. faecalis* EF67230 (g-i) for 48 h. The images showed

different magnifications as indicated on the top. The scale bar (black) represents 100 µm (a, d, g), 10 µm (b, e, h,), and 1 µm (c, f, i).

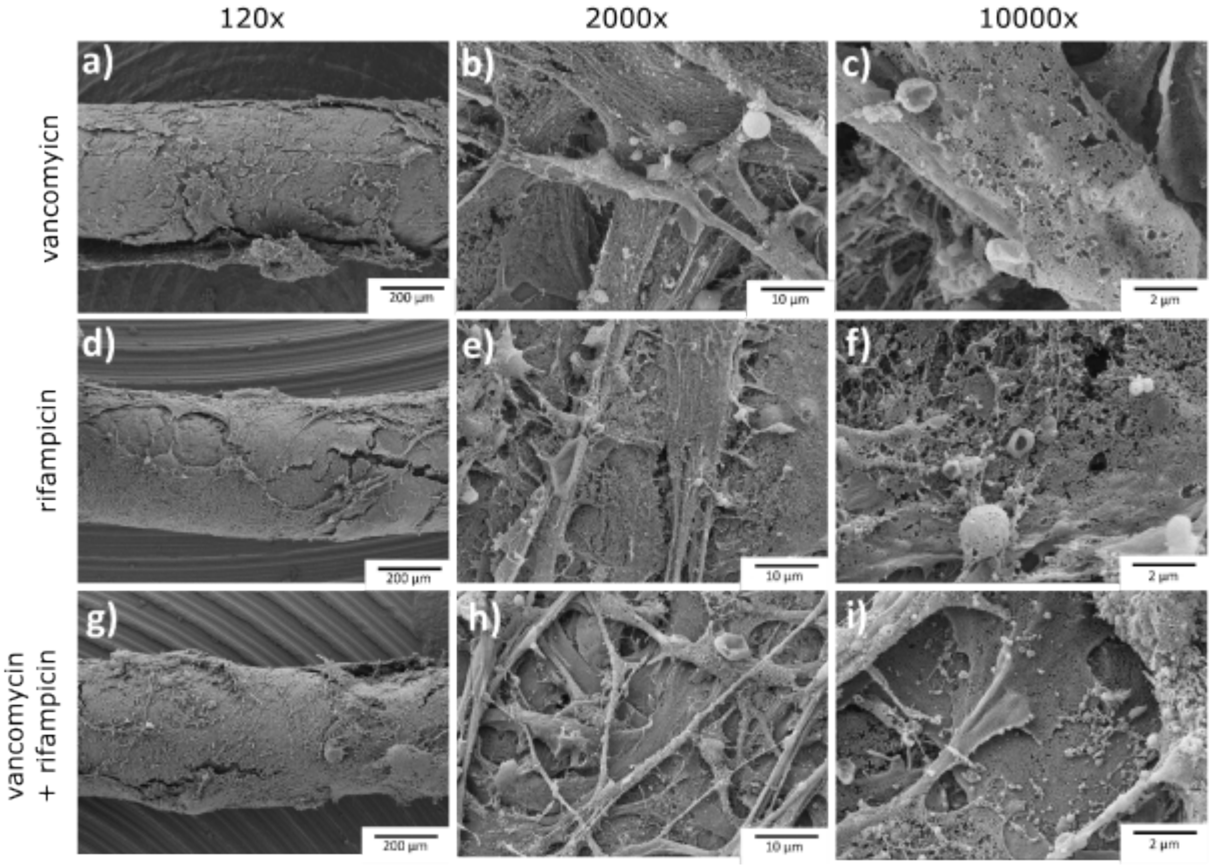


Figure S3 The post antibiotic treatment SEM analysis of *S. aureus* SA14073 biofilms formed on implants with PF methodology. The biofilm treated with 20 mg/L vancomycin (a-c), 5 mg/L rifampicin (d-f) and combination of vancomycin and rifampicin (g-i). The larvae were injected with *S. aureus* SA14073 pre-infected implant and incubated, post 48 h incubation, larvae were treated with antibiotics. After 24 h of treatment, implants were taken out of the larvae, washed with 1 x PBS, fixed and SEM was performed. The images showed different magnifications as indicated on the panels. The scale bar (black) represents 200 µm (a, d, g), 10 µm (b, e, h) and 2 µm (c, f, i).

**Table S3** Statistical analysis and significant different groups of the study.

| **Target groups** | **Significance**<br>p<0.05 (*), p<0.01 (**), p<0.001 (***), p<0.0001(****) |
|---|---|
| **Figure 2: Statistical analysis using Ordinary one-way ANOVA** | |
| SA14073 ePTFE vs. SA29552 ePTFE | * |
| SA14073 ePTFE vs. SA31685 ePTFE | * |
| SA29552 ePTFE vs. SA14073 PU | * |
| SA31685 ePTFE vs. SA14073 PU | ** |

| | |
|---|---|
| SA31685 ePTFE vs. SA29552 PU | * |
| **Figure 4: Statistical analysis using Log-rank (Mantel-Cox) test** | |
| *SA*14073 $10^7$ CFU/larvae | **** |
| *SA*29552 $10^6$ CFU/larvae | * |
| *SA*29552 $10^7$ CFU/larvae | **** |
| *SA*31685 $10^7$ CFU/larvae | **** |
| EF1653 $10^6$ CFU/larvae | * |
| EF1653 $10^7$ CFU/larvae | **** |
| EF9367 $10^6$ CFU/larvae | * |
| EF9367 $10^7$ CFU/larvae | **** |
| EF67230 $10^5$ CFU/larvae | ** |
| EF67230 $10^6$ CFU/larvae | **** |
| EF67230 $10^7$ CFU/larvae | **** |
| **Figure 5: Statistical analysis using Log-rank (Mantel-Cox) test** | |
| PBS vs SA14073 IL | ** |
| PBS vs SA14073 PF | ** |
| PBS vs SA29552 IL | *** |
| SA29552 IL vs SA29552 PF | * |
| PBS vs SA1653 IL | * |
| PBS vs SA9367 IL | **** |
| PBS vs EF67230 in vivo | **** |
| PBS vs EF67230 PF | **** |
| EF9367 IL vs EF9367 PF | **** |
| **Figure 8: Statistical analysis using Kruskal-Wallis test** | |
| 48 h PF biofilm vs. rifa 5 mg/L (SA14073) | ** |
| 48 h PF biofilm vs. vanco 20 mg/L + rifa 5 mg/L (SA14073) | *** |
| 48 h PF biofilm vs. rifa 5 mg/L (SA29552) | * |
| 48 h PF biofilm vs. vanco 20 mg/L + rifa 5 mg/L (SA29552) | ** |
| 48 h PFbiofilm vs. rifa 5 mg/L (SA31685) | ** |
| 48 h PF biofilm vs. vanco 20 mg/L + rifa 5 mg/L (SA31685) | * |
| 48 h IL biofilm vs.rRifa 5 mg/L (SA29552) | * |

| | |
|---|---|
| 48h IL biofilm vs. vanco 20 mg/L + rifa 5 mg/L (SA29552) | ** |
| 48 h IL biofilm vs. Rifa 5 mg/L (SA31685) | * |
| 48 h IL biofilm vs. vanco 20 mg/L + rifa 5 mg/L (SA31685) | * |
